# gNOMO2: a comprehensive and modular pipeline for integrated multi-omics analyses of microbiomes

**DOI:** 10.1101/2023.12.03.569767

**Authors:** Muzaffer Arıkan, Thilo Muth

**Affiliations:** Regenerative and Restorative Medicine Research Center (REMER), Research Institute for Health Sciences and Technologies (SABITA), Istanbul Medipol University, Istanbul, Türkiye; Department of Medical Biology, Faculty of Medicine, Istanbul Medipol University, Istanbul, Türkiye; Domain Data Competence Center (MF 2), Robert Koch Institute (RKI), Berlin, Germany

**Keywords:** microbiome, multi-omics, data integration, amplicon sequencing, metagenomics, metatranscriptomics, metaproteomics

## Abstract

**Background:** Over the past few years, the rise of omics technologies has offered an exceptional chance to gain a deeper insight into the structural and functional characteristics of microbial communities. As a result, there is a growing demand for user friendly, reproducible, and versatile bioinformatic tools that can effectively harness multi-omics data to offer a holistic understanding of microbiomes. Previously, we introduced gNOMO, a bioinformatic pipeline specifically tailored to analyze microbiome multi-omics data in an integrative manner. In response to the evolving demands within the microbiome field and the growing necessity for integrated multi-omics data analysis, we have implemented substantial enhancements to the gNOMO pipeline.

**Results:** Here, we present gNOMO2, a comprehensive and modular pipeline that can seamlessly manage various omics combinations, ranging from two to four distinct omics data types including 16S rRNA gene amplicon sequencing, metagenomics, metatranscriptomics, and metaproteomics. Furthermore, gNOMO2 features a specialized module for processing 16S rRNA gene amplicon sequencing data to create a protein database suitable for metaproteomics investigations. Moreover, it incorporates new differential abundance, integration and visualization approaches, all aimed at providing a more comprehensive toolkit and insightful analysis of microbiomes. The functionality of these new features is showcased through the use of four microbiome multi-omics datasets encompassing various ecosystems and omics combinations. gNOMO2 not only replicated most of the primary findings from these studies but also offered further valuable perspectives.

**Conclusions:** gNOMO2 enables the thorough integration of taxonomic and functional analyses in microbiome multi-omics data, opening up avenues for novel insights in the field of both host associated and free-living microbiome research. gNOMO2 is available freely at https://github.com/muzafferarikan/gNOMO2.

## Background

Microbiomes play pivotal roles in shaping the environments they inhabit such as influencing host health and disease (1), and contributing to the overall diversity of life on Earth (2). The comprehensive understanding of microbial communities and their impact on human health, ecosystems, and numerous other domains has become an increasingly prominent field of investigation (3).

Over the past decade, there has been a substantial increase in various omics data types generated from various microbiomes due to the development of novel techniques and reduced experimental costs (4, 5). Hence, the multi-omics approach has emerged as a powerful strategy to elucidate the functional potential of microbiomes, going beyond taxonomic profiling to decipher the molecular mechanisms (6–8). The metabolic pathways, ecological interactions, and adaptive responses of microbial communities can be uncovered by integrating multiple omics data (9). Such a comprehensive perspective is invaluable for potential implications in diverse fields, such as human health, agriculture, and environmental conservation.

To unravel the complex web of interactions within microbiomes and extract meaningful insights from the vast amount of data generated by advanced omics technologies, the development of sophisticated analytical tools and data analysis pipelines is essential (10). Consequently, many approaches and tools have emerged to address these needs (11–15). One such pipeline, gNOMO, facilitates the integrated multi-omics analysis encompassing metagenomics (MG), metatranscriptomics (MT), and metaproteomics (MP) through the efficient generation and use of a proteogenomic database, as well as differential abundance analysis-based integration at the pathway and taxa levels (16). However, gNOMO (along with other existing multi-omics analysis tools in microbiome field) currently lacks the capability of processing 16S rRNA gene amplicon sequencing (AS) data to create a proteogenomic database while it is well known that the protein sequence database directly impacts the outcome of any MP analysis (17).

For MP, it was shown that unnecessarily large databases can lead to the exclusion of valid peptide spectrum matches (18), demanding more time and memory resources. Conversely, smaller databases carry the risk of generating false positive results that are irrelevant to the sample. In multi-omics based microbiome studies that combine MP with MG or MT, protein databases are typically generated from MG and MT data. However, for studies that integrate MP with AS, there is currently no tool available to automatically create a protein database from AS data for MP analysis. Generating an AS-based protein database can also be valuable for studies that integrate MP, MG and MT, as sequencing depth limitations may affect the detection of microbes and genes, thereby influencing MP analysis. Additionally, there is a lack of tools for conducting end-to-end integrated analysis of AS data in conjunction with MP results. Most existing multi-omics analysis tools are tailored to specific omics combinations and lack a modular architecture that can accommodate various omics combinations. Furthermore, there is a shortage of multi-omics analysis tools that incorporate multiple integration approaches and present results at different analysis stages, facilitating further investigations using other tools.

To address abovementioned needs of multi-omics data analyses in microbiome research, we have made significant improvements to the gNOMO pipeline. These enhancements encompass the following key modifications: i) We have restructured the pipeline, introducing a flexible and modular architecture which empowers gNOMO2 to seamlessly process a wide array of multi-omics data derived from microbiomes. With six independent modules, gNOMO2 can effortlessly manage a vast spectrum of omics combinations, ranging from two to four distinct omics data types, which include AS, MG, MT, and MP. ii) One of standout features of gNOMO2 is its ability to process AS data and generate a protein database suitable for MP studies. iii) Additionally, gNOMO2 incorporates three distinct approaches for integrated multi-omics analysis: proteogenomic database-based integration, differential abundance-based integration (at taxa, functional category and pathway levels), and joint visualization-based integration. These innovative approaches offer a comprehensive perspective on the microbiomes, enabling researchers to gain deeper insights into the structural and functional properties. gNOMO2 is an open-source tool and freely available at https://github.com/muzafferarikan/gNOMO2.

## Methods

### Overview of the gNOMO2 pipeline

The gNOMO2 pipeline is designed as a tool that relies on Snakemake (19), a well-established bioinformatic workflow management system. This framework guarantees scalable data analyses and the generation of consistent and reproducible output data. The pipeline incorporates a suite of software tools written in various programming languages, including R, Python, Shell and Perl, enabling the seamless execution of multi-omics analysis steps for microbiome data. The input data and program parameters in Snakemake are easily defined through a straightforward configuration file. gNOMO2 streamlines this process by automatically generating the configuration file from the provided input data along with default parameters.

To enhance user experience, the pipeline relies on publicly accessible tools distributed as Conda environments, simplifying the installation process for individual software components for the end user. gNOMO2 ensures result consistency and makes it user-friendly for individuals with basic bioinformatics skills to analyze multi-omics data. The pipeline accepts raw sequencing files (in fastq.gz format) for AS, MG and MT data and input MS/MS spectrum files (in mgf format) for MP data.

The original gNOMO accepts MG, MT and MP data as input and generates results for differential abundance analysis in each omics layer. It also constructs a protein database using MG and MT data and performs both differential abundance and pathway level integrated analyses (Figure 1A). In contrast, gNOMO2 pipeline comprises six modules that facilitate direct analysis of various omics combinations. Each module includes pre-processing, analysis of each omics dataset, data integration and visualization steps (Figure 1B).

**Figure 1.**
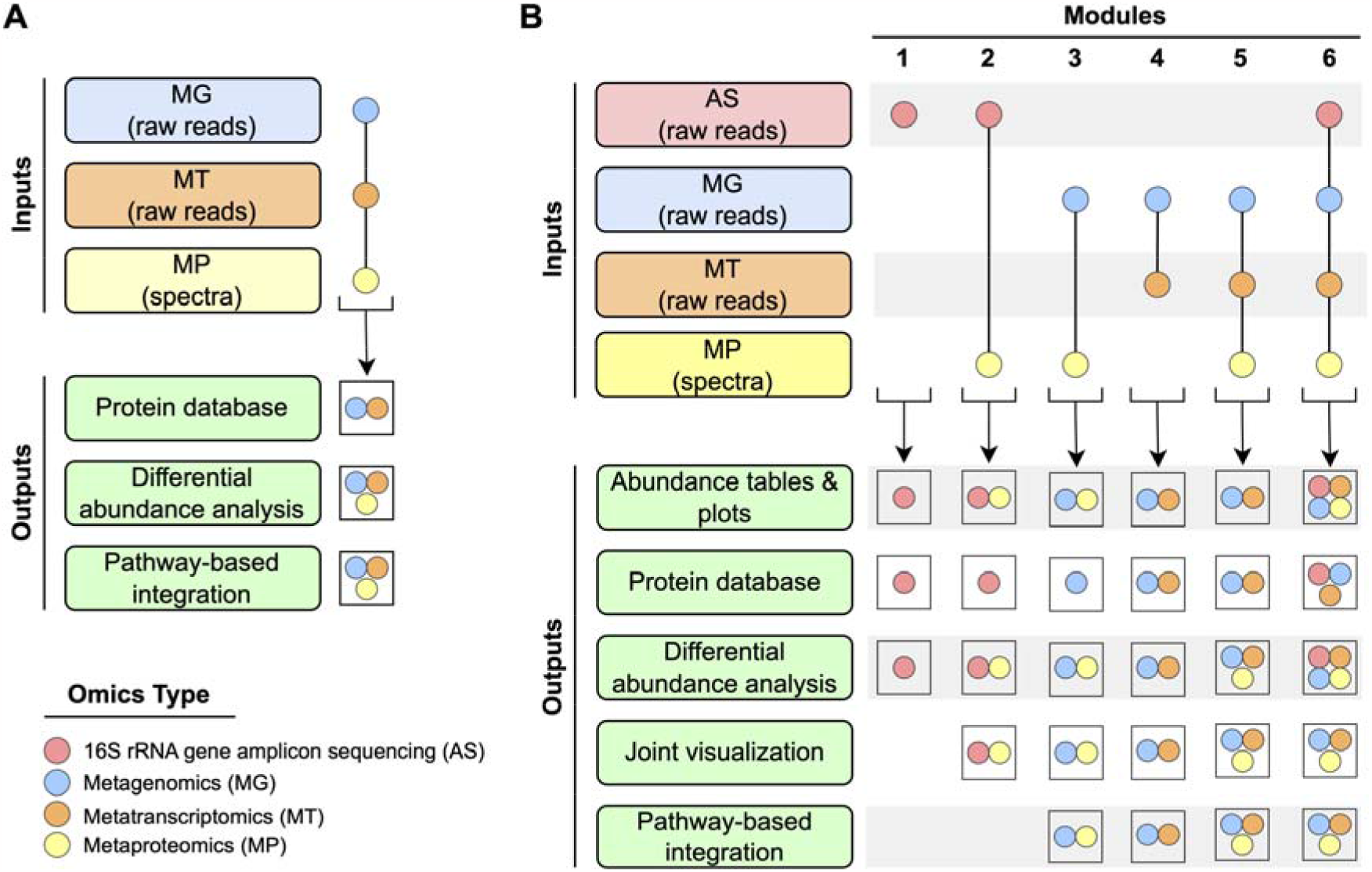
Overview of gNOMO and gNOMO2 pipelines. (A) gNOMO accepts MG, MT and MP data as input, providing differential abundance analysis results for each omics layer. It also generates a protein database using MG and MT data and performs a pathway-level integrated analysis. (B) gNOMO2 comprises six modules, each tailored for specific omics data. Module 1 accepts 16S rRNA gene amplicon sequencing data (AS) as input and generates a protein database suitable for metaproteomics studies, a taxa abundance plot and a phyloseq object that can be used for downstream analysis in other microbiome tools. Modules 2 to 6 handle different combinations of AS, MG, MT, and MP data, creating omics-specific protein databases, abundance tables, plots, differential abundance analysis results, and pathway-level integration analysis results.

#### Module 1: Processing AS data and generating a protein database for MP analysis

Module 1 is designed to process raw AS data in both paired-end and single-end formats, providing a directly usable protein database for MP data analysis. The first step in this module involves using Trimmomatic (20) to remove sequencing adapters, low quality bases from raw reads and reads that are too short (default minimum length > 25 bp). The quality of both raw and trimmed reads is assessed using FastQC (21), and analysis results for all samples are summarized using MultiQC (22). If the data is in paired-end format, the quality controlled reads are merged using FLASH (23). Subsequently, DADA2 (24) is used in conjunction with the SILVA database (25) to obtain an amplicon sequence variant (ASV) abundance table and taxonomy assignments for each ASV. After determining the user defined top n most abundant taxa at a user defined taxonomic level, protein sequences of all complete genomes for these taxa are downloaded from the National Center for Biotechnology Information (NCBI) database using the ncbi-genome-download (https://github.com/kblin/ncbi-genome-download) tool. All downloaded sequences are merged, cleaned and redundancies are then removed using SeqKit (26). Importantly, for host associated microbiome samples, the user can define host species name in a configuration file. Host protein sequences are then included in the final protein database together with microbial proteins. Module 1 allows researchers to construct a comprehensive protein database, either from their own AS datasets or publicly available ones. Furthermore, this module creates a phyloseq (27) object containing an abundance table, a taxonomy table and additional metadata. This enables ongoing microbiome analysis using other analysis tools. In addition, an abundance plot is automatically generated to assess the abundance distribution of the top ‘n’ taxa as defined by the user.

#### Module 2: Integrated multi-omics analysis of AS and MP data

Module 2 accepts raw paired-end and single-end AS and MP data as inputs. The AS data undergoes the processing steps described in Module 1. The generated AS-based protein database is then used for the database search algorithm MS-GF+ (28) to identify peptides in the raw MP data. A peptide abundance table is subsequently created by aggregating results from individual samples. Taxonomy and enzyme commission (EC) assignments for the identified peptides are carried out using Pyteomics (29) and Unipept (30). MaAsLin2 (31) is employed to determine differentially abundant taxa based on both AS and MP data. Furthermore, a joint visualization of MP and AS results is performed using the combi R package (32). The final outputs include AS and MP based abundance tables, results from differential abundance analysis, and joint visualization analysis results.

#### Module 3: Integrated multi-omics analysis of MG and MP data

Module 3 is designed to handle raw paired-end MG and MP data. MP data is processed as outlined in Module 2. MG raw reads are quality checked and cleaned using Trimmomatic (20), followed by merging with FLASH (23). The quality of both raw and trimmed reads is assessed using FastQC (21), and analysis results for all samples are summarized using MultiQC (22). Cleaned and merged reads are then mapped to the NCBI non-redundant (nr) database using Kaiju (33), which generates taxonomic classification results. In parallel, clean reads are also used for assembly with metaSPAdes(34) and contigs are classified as eukaryotic and prokaryotic using EukRep (35). Proteins within the prokaryotic contigs are predicted using Prodigal (36) while Augustus (37) is used for proteins within eukaryotic contigs. Then, functional annotation of these predicted proteins is carried out using EggNOG (38) to obtain KEGG Orthology (KO) identifiers, while InterProScan (39) is employed for TIGRFAM (40) functional annotation. The final outputs of Module 3 for both MG and MP results include taxonomic composition plots, taxa and functional annotation based differential abundance analysis, joint visualization of omics layers as described in Module 2 and metadata parameters. Moreover, pathway level integrated analysis is conducted using Pathview (41) package. The Pathview plots represent the log2 ratio of the means of the different user defined conditions at different omics levels after a fold change normalization. These log2 ratios are calculated and compared through shared enzyme and KEGG ids. Coverages for gene sequences of each predicted protein by MG are calculated using BBMap (42).

#### Module 4: Integrated multi-omics analysis of MG and MT data

Module 4 is designed to handle raw paired-end MG and both paired-end and single-end MT data. MG data follows the processing steps outlined in Module 3. For MT data, a similar workflow is employed, with the exception that a *de novo* assembly step is conducted using rnaSPAdes (43) in place of metaSPAdes. The final outputs of Module 4 include an MG&MT-based proteogenomic database, taxonomic and functional annotation based differential abundance analysis results for both omics levels, abundance tables and plots, joint visualization of omics layers as described in Module 2, metadata parameters, pathway-level integrated analysis results as outlined in Module 3.

#### Module 5: Integrated multi-omics analysis of MG, MT and MP data

Module 5 accepts raw paired end MG, both paired-end and single-end MT and MP data. MG and MT data follow the processing steps outlined in Module 4 while MP data is processed as described in Module 2. The final outputs of Module 5 include a MG&MT-based proteogenomic database, taxonomic and functional annotation based differential abundance analysis results for three omics levels, abundance tables and plots, joint visualization of omics layers and metadata parameters and pathway level integrated analysis results as outlined in Module 3.

#### Module 6: Integrated multi-omics analysis of AS, MG, MT and MP data

Module 6 accepts both paired-end and single-end AS and MT data, paired end MG, and MP data. MG, MT and MP data follow the processing steps outlined in Module 5. However, the final outputs of Module 6 include a proteogenomic database which is generated by combining AS, MG and MT based downloaded/predicted protein sequences, taxonomic and functional annotation based differential abundance analysis results for four omics levels, abundance tables and plots, joint visualization of omics layers, metadata parameters and pathway level integrated analysis results as outlined in Module 3.

## Results and discussion

To illustrate the utility of gNOMO2, we re-analyzed samples from four previously published microbiome studies involving various multi-omics combinations, using the respective publicly available datasets.

### Analyzing the association of saliva content with oral cancer

Saliva is a complex biofluid that comprises various components, including DNA, RNA, proteins, metabolites, and microbiota. As a result, it is considered as a promising source of relevant biomarkers for a variety of diseases (44). Granato *et al*. (2021) combined AS and MP analyses to investigate the association between saliva content and oral cancer (45). The study suggests that oral microbiota and their protein abundance have potential diagnosis and prognosis value for oral cancer patients. Here, we showcase how Modules 1 (AS) 2 (AS and MP) of gNOMO2 can be used to efficiently reproduce the findings.

The AS data was obtained from NCBI SRA under BioProject identifier PRJNA700849 while MP data was retrieved from PRIDE under accession number PXD022859. The dataset included saliva samples from 8 healthy controls and 15 oral cancer patients. To streamline downstream analyses, we merged triplicates of AS samples and used cell debris MP samples for all analyses. The taxonomic composition results based on AS data across samples, as generated by gNOMO2, were consistent with the reported results, demonstrating similar abundance distributions and the presence of the same most abundant genera (Figure 2A). In their study, Granato *et al*. (2021) constructed a protein database containing 1,160,275 protein sequences from the 12 most abundant bacterial genera and human. We applied the same parameters in gNOMO2 to achieve comparable results, with setting such as taxa_level: Genus, top_n: 12 and host: Homo sapiens. gNOMO2 automatically generated a protein database from AS data by determining the 12 most abundant bacterial genera. It then retrieved all protein sequences from 1,992 genomes belonging these bacterial genera, along with human host proteins, resulting in a total of 1,240,988 protein sequences. The discrepancy in the number of protein sequences between the generated protein databases may be attributed to variations in analysis timing and database differences. Granato *et al*. (2021) used the HOMD, a specific database used for oral microbiome studies while gNOMO2 uses the NCBI database, intended to target all microbiome study types.

**Figure 2.**
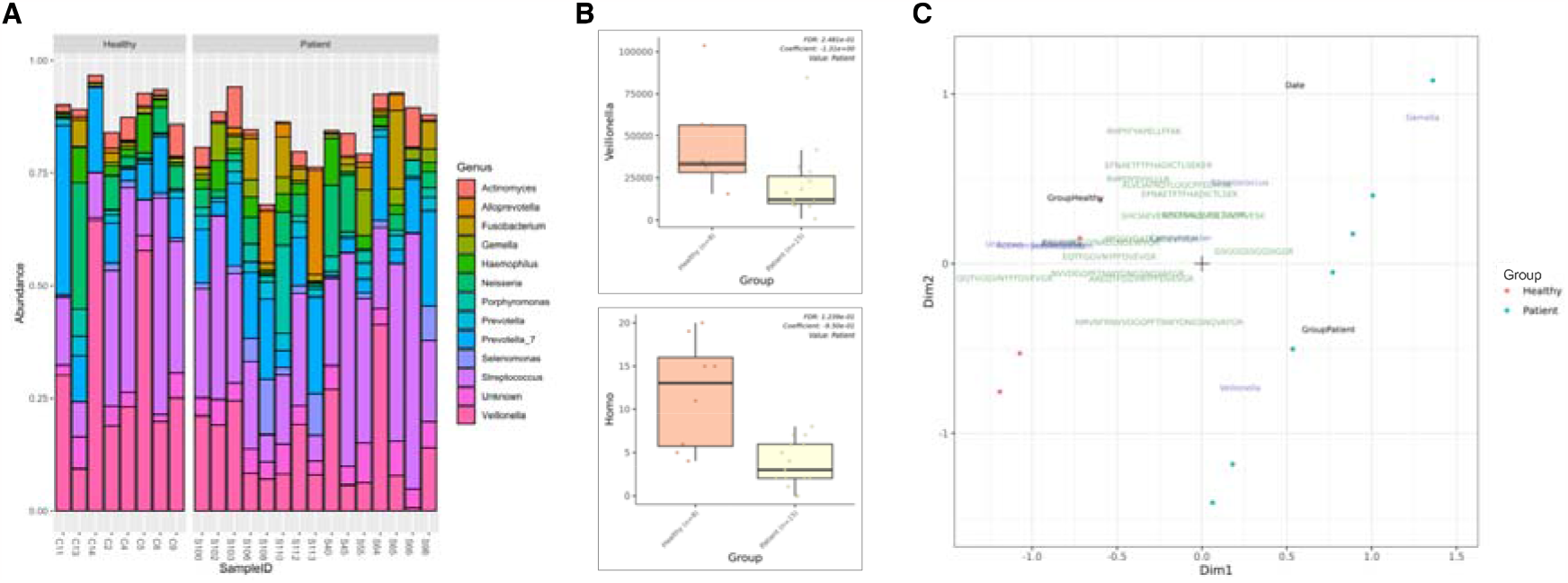
Overview of gNOMO2 results for the Granato *et al*. (2021) study. (A) Representation of the ten most prevalent genera in saliva microbiota samples. AS-based representations of salivary microbiota composition across samples, highlighting the ten most common bacterial genera. Each bar indicates the relative abundance distribution for a sample. (B) Abundance distribution of differentially abundant taxa across study groups, presented separately for AS (upper) and MP (lower) data. (C) Joint visualization-based integration results for AS, MP and metadata. Blue labels represent taxa, green labels represent peptides, and black labels represent metadata columns. Patient samples are marked with blue dots, while healthy samples are marked with red dots.

Within gNOMO2 users can also perform differential abundance analysis at both omics levels, yielding statistical test results and plots for differential abundant taxa. For instance, we presented one of differential taxa from AS-based (Figure 2B, upper) and MP-based results (Figure 2B, lower). AS-based differential abundance analysis showed a decrease in the abundance of *Veillonella* associated with oral cancer (Figure 2B, upper), which corresponds to a key finding in the Granato *et al*. (2021) study and previous studies (46). Interestingly, gNOMO2 detected a reduction in the abundance of peptides classified as Homo in oral cancer patients (Figure 2B, lower) while the original study did not report any statistically significant changes. This divergence may result from differences in analysis approaches, as gNOMO2 employs a peptide-based taxonomy by Unipept and MaAslin2 for differential abundance analysis instead of a protein-based approach. Furthermore, it is important to note that we did not account for other covariates that may affect the results.

Finally, gNOMO2 generates a joint visualization plot for AS, MP and metadata (Figure 2C). This plot confirms the association of *Veillonella* based on AS with oral health status based on AS data and additionally reveals associations between some detected peptides and the oral health status of the participants. Notably, InterPro entries assigned to these peptides included human albumin proteins, which were previously reported to be associated with oral cancer (47, 48).

### Exploring potential and active functions within the human gut microbiota

The human gut microbiota is widely recognized for its important roles in both health and disease. A comprehensive understanding of both potential and active features can provide valuable insights into the mechanisms governing various physiological processes and pathologies, ultimately leading to more effective strategies for maintaining and improving human well-being.

Tanca *et al*. (2017) employed MG and MP to explore the potential and active functions in the gut microbiota of a healthy human cohort (49). Here, we used Module 3 (MG and MP) of gNOMO2 to efficiently re-analyze the multi-omics data from their study. The MG data was obtained from the NCBI SRA under BioProject identifier PRJEB19090, while the MP data was retrieved from PRIDE under accession number PXD005780. The dataset included gut microbiota samples from 6 males and 8 females.

We employed gNOMO2 to investigate potential differences between male and female participants. Taxonomic composition results based on MG and MP data, as generated by gNOMO2, exhibited a significant overlap with the findings of Tanca *et al*. (2017), particularly concerning the most abundant genera (Figure 3A, upper). MG-based differential abundance analysis, using default parameters, indicated a notably higher abundance of *Legionella* in females. Nevertheless, it is important to approach this finding with caution, given that *Legionella* is a bacterial genus typically associated with water and soil environments, often considered a potential source of contamination in human microbiome studies (50).

**Figure 3.**
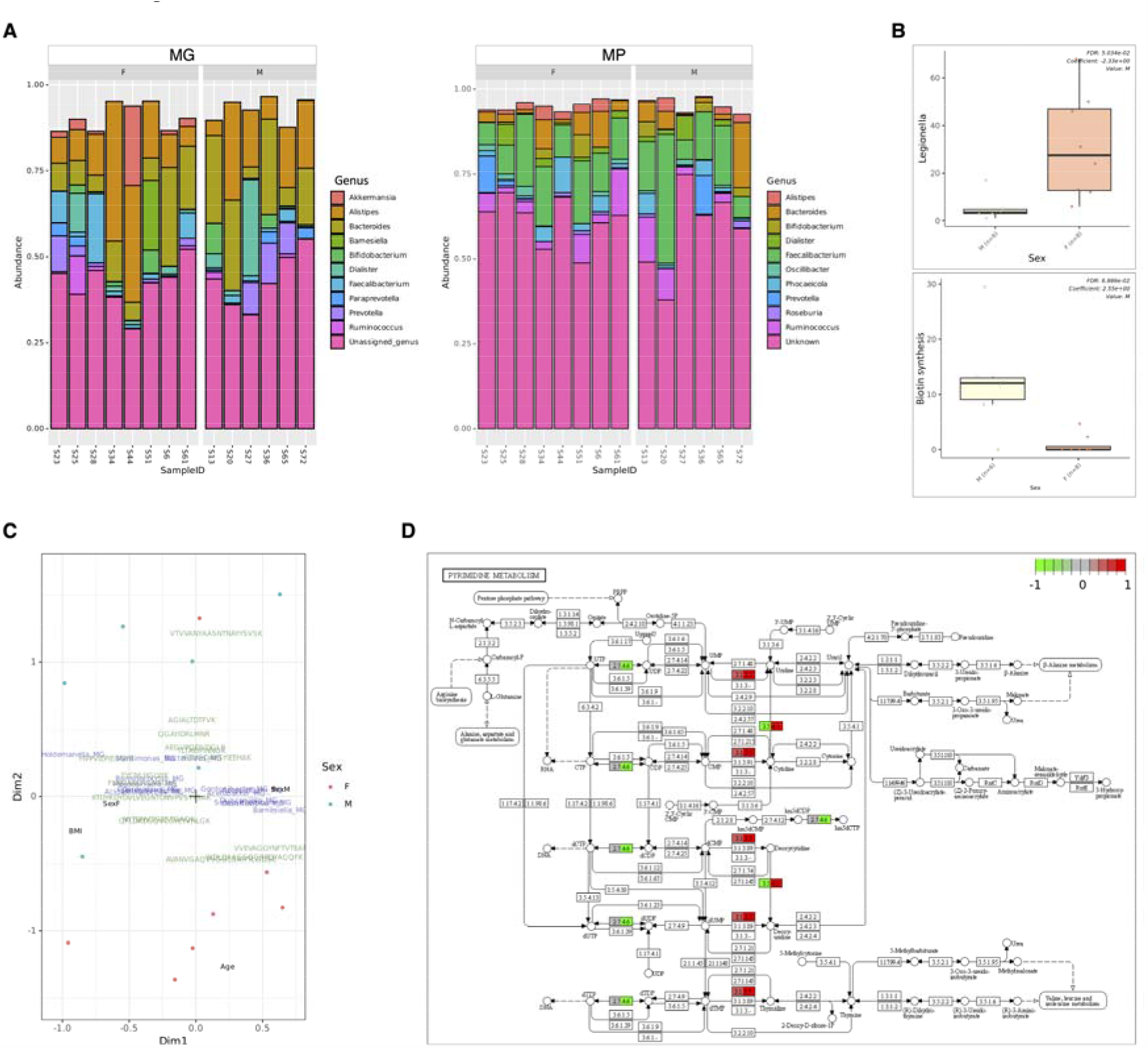
Overview of gNOMO2 results for Tanca *et al*. (2017) study. (A) Representation of the ten most prevalent genera in gut microbiota samples, as shown by MG and MP. The left side illustrates the ten most common bacterial genera based on MG data, while the right side represents MP-based findings. Each bar represents relative abundance distribution for a sample. (B) Abundance distribution of differentially abundant taxa across study groups, separately for MG (upper) and MP (lower) data. (C) Joint visualization-based integration results for MG, MP and metadata. Blue labels represent taxa, green labels show peptides and black labels represent metadata columns. Male samples are marked with blue dots, while female samples are marked with red dots. (D) Pathway level integration results, demonstrating the relationship across different omics levels. The findings from MG and MP are illustrated separately as split nodes on the left and right, respectively.

Functional annotations derived from TIGRFAM for the differential abundance analysis indicated a reduction in biotin synthesis (Figure 3B, lower). The joint visualization plot depicted both MG and MP features along with covariates such as BMI, age and sex (Figure 3C). In our pathway-level integration analysis, we illustrated the components of pyrimidine metabolism and how variations in their abundance can be observed among study groups across different omics levels (Figure 3D). As a case in point, cytidine deaminase (EC 3.5.4.5) displayed an elevated abundance in females at the MG level (colored red, left), while its abundance decreased at the MP level (colored green, right). This discrepancy suggests an increase in the abundance of taxa carrying the corresponding gene but a lower expression of the protein. Again, this highlights the significance of adopting a multi-omics perspective when drawing conclusions in microbiome studies.

### Investigating the role of microbiota of the Maasdam cheese during ripening

The microbiota present in cheese plays a crucial role in the maturation and development of its distinctive flavor, making it a pivotal aspect for the cheese industry. Duru *et al*. (2018) combined MG and MT to track shifts in both taxonomic compositions and gene expressions of Swiss-type Maasdam cheese microbiota during the ripening process (51). Here, we used Module 4 (MG and MT) of gNOMO2 to efficiently re-analyze multi-omics data from their research.

MG and MT data were retrieved from the EBI ENA under BioProject identifier PRJEB23938. The dataset comprised three samples from day 12 and three samples from day 37 day of the ripening process.

We employed gNOMO2 to investigate potential differences between different stages of ripening process. Taxonomic composition results generated by gNOMO2 based on MG and MT data showed that *Lactococcus, Lactobacillus* and *Propionibacterium* were three most abundant genera across samples (Figure 4A), in consistent with the findings of Duru *et al*. (2018). Differential abundance analyses revealed significantly higher relative abundance of *Propionibacterium*, the main bacterial genera responsible for propionate metabolism in the Maasdam cheese, in cold ripening samples in both MG and MT levels (Figure 4B) which is also well aligning with the results of the original study.

**Figure 4.**
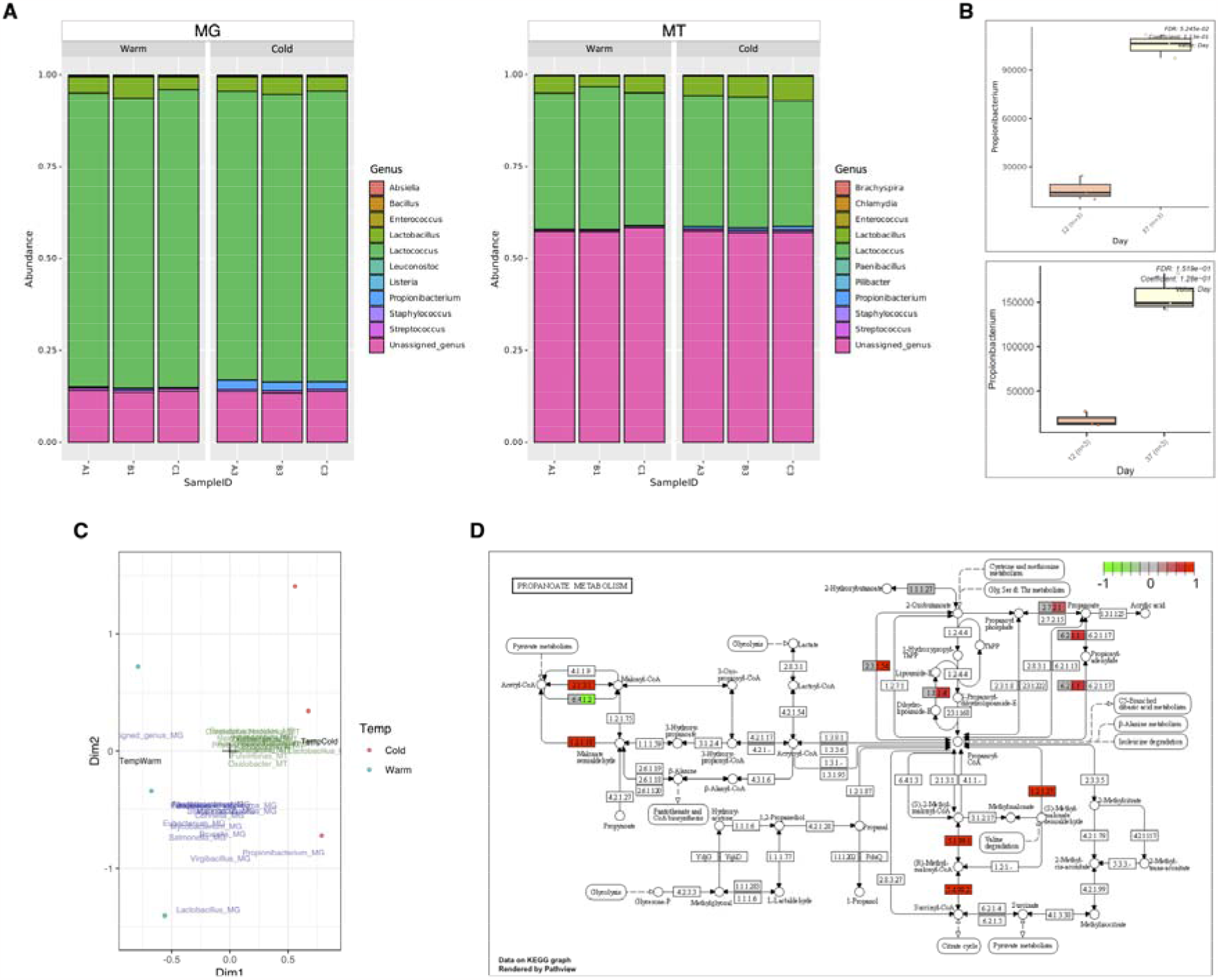
Overview of gNOMO2 results for the Duru *et al*. (2018) study. (A) Representation of the 10 most common genera in cheese microbiota samples. MG- and MT-based overview of gut microbiota composition across samples. The 10 most common bacterial genera in cheese microbiota samples are shown for MG (left) and MT (right). Each bar represents relative abundance distribution for a sample. (B) Abundance distribution of differentially abundant taxa across study groups by MG (upper) and MT (lower). (C) Joint visualization-based integration results for MG, MT and metadata. (D) Pathway level integration results, demonstrating the relationship across different omics levels. The findings from MG and MT are illustrated separately as split nodes on the left and right, respectively.

The joint visualization plot depicted both MG and MT features along with the ripening types (Figure 4C). In our exploration of pathway-level integration, we depicted the elements of propionate metabolism and highlighted how fluctuations in their abundance varied across study groups at MG and MT levels (Figure 4D). Notably, genes related to propionate production exhibited higher abundance in cold ripening samples (day 37) compared to warm ripening ones (day 12) at the MT level (colored red, right), while their levels were not significantly different at the MG level (colored gray, left). As a result, we did not observe a decrease in expression of genes responsible for propionate production, contrary to findings in the original study. This discrepancy may originate from methodological differences between the studies. The gNOMO2 pipeline compares the expression of propionate production genes against total gene expression, whereas Duru *et al*. (2018) study compared these genes against the overall expression of the *Propionibacterium* genome obtained in their research. Consequently, the relative expression of these genes might appear higher when assessed against all genes but lower when measured against only *Propionibacterium* genes. To validate this, we conducted comparisons using the *Propionibacterium* genome from the original study in the gNOMO2 pipeline for gene expression levels. Changing the denominator from all genes to *Propionibacterium* genes yielded results consistent with the original study. This underscores the importance of accurately interpreting analysis outcomes based on the structure of the analytical pipeline.

### Determining microbiome dynamics in a wastewater treatment plant

Characterization of microbial communities across various meta-omics layers offers important insights into their potential traits and functionalities. Herold et al. (2020) utilized MG, MT, MP, and metabolomics to explore the responses of microbial populations in a biological wastewater treatment plant to disturbances. In our study, we demonstrate how Modules 5 (MG, MT, and MP) and 6 (AS, MG, MT, and MP) of gNOMO2 effectively replicate some of their findings using a subset of the samples.

We obtained AS, MG, and MT sequencing data from EBI ENA (BioProject identifier PRJNA230567) and MP data from PRIDE (accession number PXD013655). To investigate seasonal variations reported by Herold et al. (2020), we selected samples showcasing the most distinct differences between summer and winter seasons, encompassing five samples from each. Additionally, we incorporated 10 AS samples previously collected from the same wastewater treatment plant by the same research group to assess Module 6.

Our analysis, performed using gNOMO2, revealed taxonomic composition results (AS, MG, MT, and MP data) that partially aligned with Herold et al.’s findings (Figure 5A). However, unlike the original study, we did not observe pronounced compositional changes in winter samples (Figure 5A). This discrepancy may be attributed to differing approaches in taxonomic composition analysis as Herold et al. utilized taxonomic assignments of a subset of metagenome assembled genomes, while gNOMO2 employs Kaiju for direct taxonomic classification of reads.

**Figure 5.**
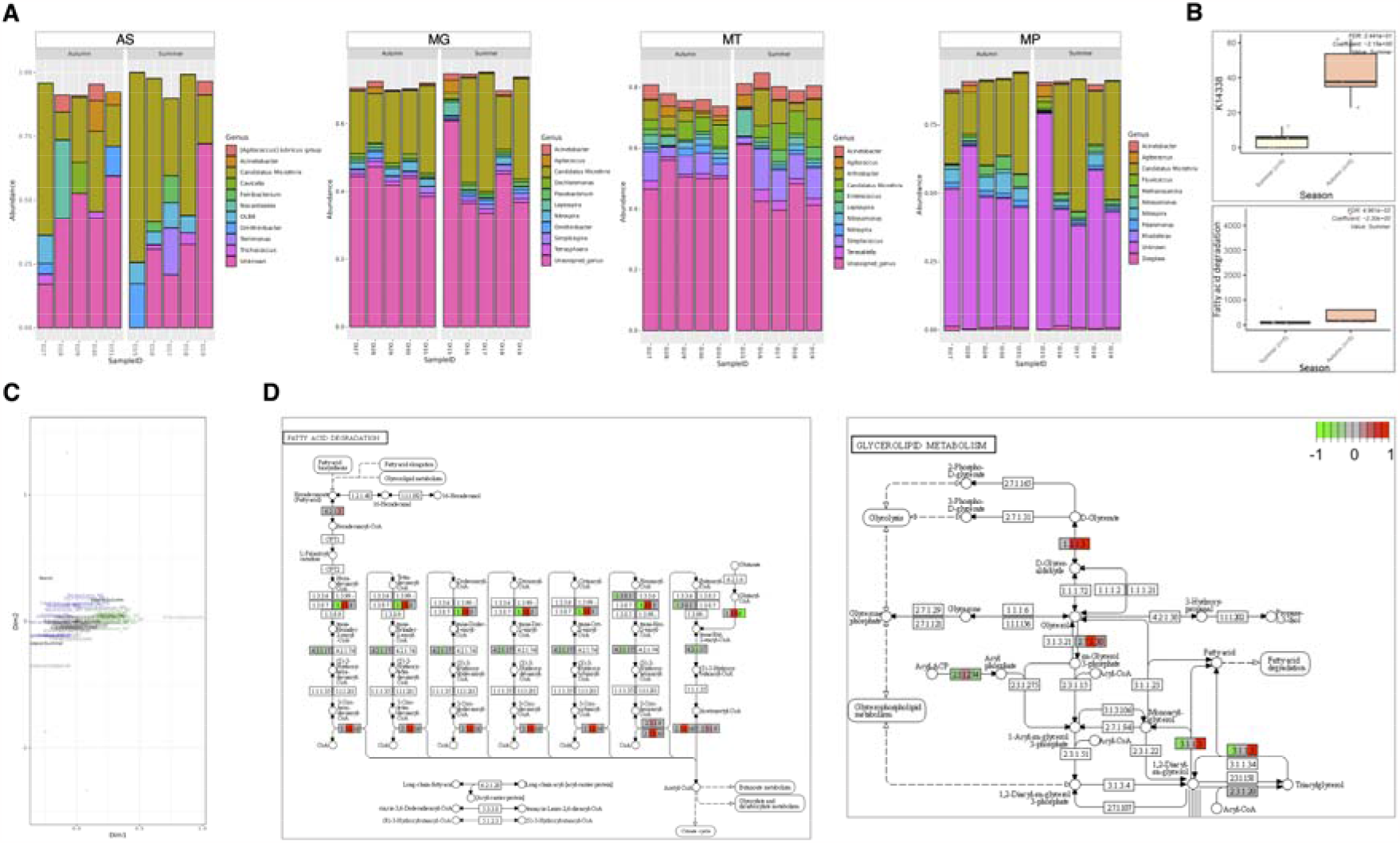
Overview of gNOMO2 results for the Herold *et al*. (2020) study. (A) Representation of the 10 most common genera in wastewater microbiota samples. MG-, MT- and MP-based overview of gut microbiota composition across samples. The 10 most common bacterial genera in wastewater microbiota samples by MG (left), MT (middle) and MP (right). Each bar represents relative abundance distribution for a sample. (B) Abundance distribution of differentially abundant taxa across study groups by MG (upper), MT (middle) and MP (lower). (C) Joint visualization-based integration results for MG, MT, MP and metadata. (D) Pathway level integration results, demonstrating the relationship across different omics levels. The findings from MG, MT, and MP are illustrated separately as split nodes on the left, middle, and right, respectively.

While gNOMO2 did not detect differentially abundant taxa between seasons across MG, MT, and MP layers, our TIGRFAM and KEGG pathway-based analyses indicated an elevation in fatty acid degradation at the MT level (Figure 5B), aligning with the original study. The joint-visualization plot highlighted MG, MT and MP features along with covariates (Figure 5C). As a case point, the plot revealed the association of *Tetrasphaera* with autumn which has been reported in previous studies to be associated with sludge bulking that frequently occurs in wastewater treatment plants (52, 53).

In our pathway-level integration analysis (Figure 5D), we illustrated variations in the components of fatty acid degradation and glycerolipid metabolism among study groups across different omics levels. Specifically, gNOMO2 showcased an increase in fatty acid degradation at the MT level while detecting an elevation in glycerolipid metabolism at both MT and MP levels, as indicated and discussed in detail in the original paper.

When AS data were integrated using Module 6, gNOMO2 constructed a proteogenomic database comprising 4,959,677 proteins, incorporating 859,729 non-redundant proteins derived from the top 10 most abundant genera identified in the AS analysis, in addition to the 4,025,111 proteins obtained from MG and MT analyses. Interestingly, this integration resulted a slight decrease in the number of detected unique peptides (∼2%), indicating the importance of database size optimization in the multi-omics studies including MP. The inclusion of AS data did not alter the other outcomes derived from the MP data analysis.

## Conclusions

gNOMO2 stands as a versatile and modular bioinformatic pipeline designed for integrated multi-omics analyses of AS, MG, MT, and MP data in a reproducible fashion. Our open-source tool efficiently employs techniques that process raw data and generates summary tables and figures with just a single, straightforward command. gNOMO2 encompasses preprocessing, genome mapping, assembly, protein predictions, taxonomic and functional annotations, proteogenomic database generation and differential abundance analysis steps for each omics layer. Furthermore, gNOMO2 offers a holistic perspective through integrated visualization of omics layers and facilitates pathway-level integrative analysis. In addition, it includes a dedicated module for AS data processing and the automatic protein database generation for MP studies. gNOMO2 generates results that can serve as inputs for subsequent microbiome analyses using various bioinformatics tools, enhancing user flexibility throughout the process. Demonstrated efficacy of gNOMO2 with real datasets underscores it as an invaluable tool across various multi-omics combinations in microbiome research. Finally, the emphasis on reproducibility is a cornerstone of gNOMO2, as it not only streamlines the analytical process but also ensures the reliability of results by providing users with fully documented and executable workflows, enhancing the transparency and replicability in omics-driven microbiome research.

### Limitations and future directions

Despite its usefulness and effectiveness in multi-omics based microbiome research, gNOMO2 still has certain limitations. Firstly, its performance may be influenced by the quality and depth of input data, thereby necessitating potential parameter optimizations by the user. Secondly, gNOMO2 relies on existing databases for taxonomic and functional annotations which may restrict the detection of features not cataloged within these databases. Moreover, gNOMO2’s efficacy may also be influenced by the complexity of microbial communities, particularly in cases of high diversity or rare taxa, where accurate profiling may be challenging. Lastly, users should be aware that gNOMO2 assumes a certain level of computational proficiency, and while efforts have been made to enhance user-friendliness, beginners may still face a learning curve because there is no graphical user interface provided.

Future versions of gNOMO2 could address these limitations through continuous updates, improved algorithmic approaches, and increased flexibility in handling diverse omics types, datasets and microbial community structures.

## Availability and requirements

### Project name

gNOMO2

### Project home page

https://github.com/muzafferarikan/gNOMO2

### Operating system(s)

GNU/Linux

### Programming language

Python, R, Shell and Perl

### Other requirements

Conda and Snakemake are required for implementation. At least 1 TB hard drive space and 200 GB memory are recommended to run the pipeline, dependent on databases and input file sizes used.

### License

MIT

### Restrictions to use by non-academics

No

## Abbreviations

NCBI: National Center for Biotechnology Information,
ENA: European Nucleotide Archive,
MIT: Massachusetts Institute of Technology,
SRA: Sequence Read Archive.

## Acknowledgements

Muzaffer Arıkan is a recipient of the Scientific and Technological Research Council of Turkey (TUBITAK), BIDEB 2219-International Postdoctoral Research Fellowship.

## Availability of data and materials

The gNOMO2 pipeline is available at GitHub (https://github.com/muzafferarikan/gNOMO2).

## Authors’ contributions

MA and TM conceived the idea and designed the pipeline. MA implemented the pipeline, performed analyses and wrote the manuscript. TM reviewed and edited the manuscript. All authors approved the final version of the manuscript.

## Competing interests

The authors declare that they have no competing interests.

## Notes

### Competing Interest Statement

The authors have declared no competing interest.

